# Genome-wide identification of loss of heterozygosity generated by mitotic homologous recombination reveals its possible association with spatial positioning of chromosomes

**DOI:** 10.1101/2022.06.14.496055

**Authors:** Hyeonjeong Kim, Mikita Suyama

**Affiliations:** Division of Bioinformatics, Medical Institute of Bioregulation, Kyushu University, Maidashi 3-1-1, Higashi-ku, Fukuoka, 812-8582, Japan

## Abstract

Loss of heterozygosity (LOH) is a genetic alteration that results from the loss of one allele at a heterozygous locus. Some LOH events are generated by mitotic homologous recombination after monoallelic defection, then the novel homozygous locus has two copies of the normal counterpart allele. This phenomenon can serve as a source of genome diversity and is associated with various diseases. To clarify the nature of the LOH such as the frequency, genomic distribution, and inheritance pattern, we made use of whole-genome sequencing data of the three-generation CEPH/Utah family cohort, with the pedigree consisting of grandparents, parents, and offspring. We identified an average of 40.7 LOH events per individual taking advantage of 285 healthy individuals from 33 families in the cohort. On average 65% of them were classified as gonosomal-mosaicism-associated LOH, which exists in both germline and somatic cells. We also confirmed that the incidence of the LOH has little to do with the parents’ age and sex. Furthermore, through the analysis of the genomic region including the LOH, we found that the chance of the occurrence of the LOH tends to increase at the GC-rich locus and/or on the chromosome having a relatively close inter-homolog distance. We expect that these results provide significant insights into the association between genetic alteration and spatial position of chromosomes as well as the intrinsic genetic property of the LOH.

**Author Summary:** Loss of heterozygosity (LOH) is a common genetic alteration that a heterozygous locus becomes a homozygous locus. In some cases, if a monoallelic defection accompanying homologous recombination between inter-homolog occurs, it results in copy-neutral LOH having two copies of the allele. Although the LOH is potentially important in understanding pathogenesis of various diseases, its fundamental features have received scant attention. To characterize the nature of the LOH, data from single individuals and their parents are required. This is because it would be difficult to discriminate between the LOH and a normal homozygous allelic state using genomic data obtained only from a single individual. Whole-genome sequencing data of 33 CEPH/Utah families, which comprise large three-generation family units, motivated us to perform a genome-wide identification of the LOH. Using this dataset, we successfully identified the LOH and analyzed its frequency, genomic distribution, and inheritance pattern. Moreover, we revealed that the occurrence of the LOH was affected by the inter-homolog distances, which reflect the chromosome territory. Our findings pertaining to the LOH provide insight into the association between genetic alteration and spatial positioning of chromosomes.

## Introduction

Genetic alteration is the source of genome diversity and has the potential to give rise to genomic evolution. Loss of heterozygosity (LOH), one such genetic alteration, is a homozygotization that results from the loss of the heterozygous state via monoallelic defection in diploid cells. Particularly, if the monoallelic defection is repaired by mitotic homologous recombination using the normal counterpart allele, the locus becomes copy-neutral LOH [1].

The LOH having a neutral copy number has been reported to be implicated in malignancies as illustrated by Knudson’s Two-Hit Hypothesis and also known to be associated with other human diseases. The event often has adverse effects on cells by eliminating a copy of a normal allele and duplicating a copy of a mutated allele in a specific locus such as an oncogene and tumor suppressor gene. For example, myeloproliferative disorder was reported to be caused by gain-of-function mutations in *JAK2* gene followed by LOH [2]. Subsequently, studies showed that various diseases like myelodysplastic syndrome occur and deteriorate via LOH as assessed using technologies for genetic analysis, such as SNP array and next-generation sequencing [3–6].

Although the LOH is a functionally important genetic event, little is known about its nature; for example, its frequency and genomic distribution remain unclear. If we attempted to identify the LOH using genomic data obtained only from a single individual, it would be difficult to discriminate between the LOH and a normal homozygous allelic state. Therefore, data from single individuals and their parents are required to perform this type of analysis with precision. The same holds true for the identification of *de novo* mutations (DNMs), which are also difficult to distinguish from the normal heterozygous allelic state. Sasani recently conducted an investigation of the inheritance patterns of DNMs as well as identification using the whole-genome sequencing data of CEPH/Utah families [7]. Their analysis motivated us to use the same data to identify the LOH. This dataset comprises large three-generation family units consisting of grandparents, parents, and several offspring in Utah in the United States [8]. In particular, the Utah population has a traditionally high birth rate, which is a very powerful advantage in this type of analysis [7].

Here, using 285 individuals from 33 families in Utah in the United States, we identified the LOH events over the whole-genome. We then assessed the characteristics of the LOH, which is an exclusively postzygotic event, unlike DNM, by examining the correlation between the incidence of the event and the parents’ features, such as age and sex. Moreover, the LOH events were classified into two groups, those existing in the germline and somatic cells, by investigating the mode of inheritance of the LOH events. We thus confirmed that the LOH events are transmitted to progeny, showing that the LOH events are also present in the germline cells and contributes to the genetic diversity. Finally, we found that the occurrence of the LOH, which is generated by mitotic homologous recombination, might be associated with the inter-homolog distances, which reflect the chromosome territory (CT).

## Results

### Identification of LOH events that is generated by mitotic homologous recombination using single nucleotide variant (SNV) data

We investigated LOH that is generated via gene conversion by mitotic homologous recombination using 33 large families from Utah in the United States (Fig 1A). Although the original CEPH/Utah family dataset comprised 603 individuals from 33 large families who exhibited a blood relation [7], we selected 285 individuals from 33 immediate family units based on the married couples in the second-generation for this study (S1 Fig), because the second-generation individuals have data pertaining to both biological parents and offspring, thus allowing the direct identification of such LOH and examination of the inheritance of these events. To identify the LOH, first, we detected the LOH-defining SNV (L-SNV), which was considered to be included in the LOH region because it indicated the genotype of the LOH by comparing the genotype of second-generation individuals with those of their parents (first-generation individuals). Simply, we regarded the homozygous variants that indicated an impossible combination from the parents’ genotype as L-SNVs; for example, although the possible combinations resulting from parents with A/T and A/A genotypes are A/A or A/T, if the offspring’s genotype is T/T, then this SNV is deemed to be an L-SNV (Fig 1A) (see Methods for details). Subsequently, to examine the mode of the inheritance of the LOH, we tracked the transmission of the LOH of second-generation individuals to their offspring (third-generation individuals) by phasing the genotype throughout all three generations in the pedigree.

**Fig 1.**
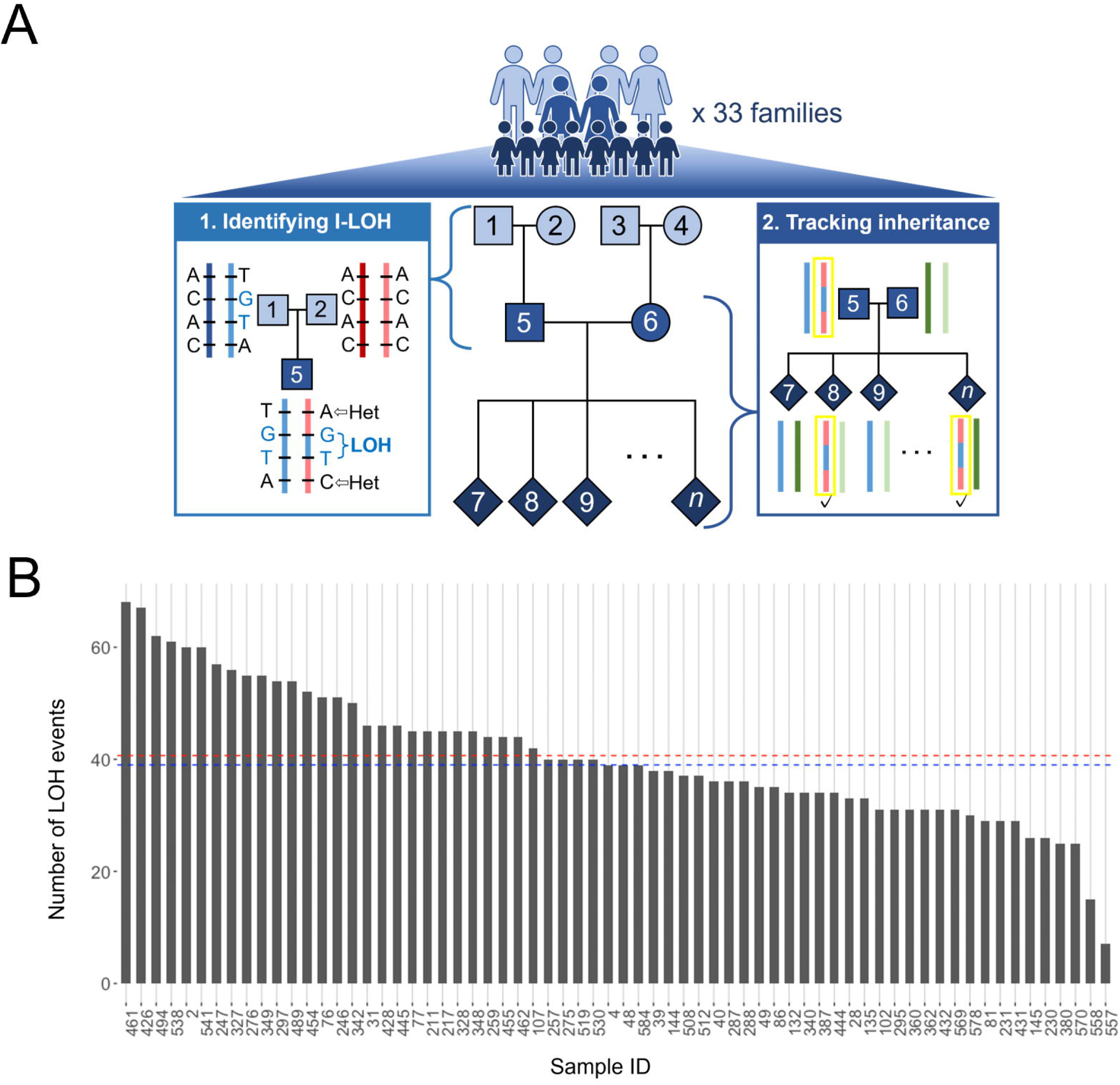
Identification of LOH events using the SNV data from the CEPH/Utah families. (A) Schematic workflow of the study. Families in the CEPH/Utah dataset consist of three generations. The method of division of a large family is described in S1 Fig. The LOH events were identified by comparing the genotype of first- and second-generation individuals. The events presented a homozygous state that originated from the genotype that existed only in one parent (blue letter in the left box). The size of LOH was restricted by the closest heterozygous variant that existed both upstream and downstream. The detailed strategy to determine the size of LOH is presented in S2 Fig. Subsequently, the mode of inheritance of the events was tracked by comparing the sequence block in the second- and third-generation individuals. (B) Total number of LOH events identified in 66 second-generation individuals. The bar graph depicts the number of events. The blue and red dashed lines indicate the median and mean numbers of the events, respectively.

We detected 4,296 L-SNVs in the entire cohort of 66 second-generation individuals from the 33 Utah families. The mean and median numbers of L-SNVs identified in an individual were 65.1 and 58.5, respectively. Then, we identified 2,684 LOH events by merging the L-SNVs depending on the heterozygous SNVs (Fig 1B). The number of L-SNVs included in an LOH was 1 to 43. Moreover, 7 (sample ID 557) to 68 (sample ID 461) LOH events were observed in each, and the mean and median numbers of events were 40.7 and 39, respectively. In addition, we estimated the minimum and maximum size of the events according to L-SNVs and heterozygous variants. The minimum size corresponds to the distance from the first L-SNV to the last L-SNV in an LOH event, and the maximum size corresponds to the distance between two nucleotides just before the first heterozygous variant upstream and downstream from an LOH event (S2 Fig). Based on this criterion, the length range of the minimum size was 1 to 155,999 bp, with a median size of 1 bp. In turn, the length range of the maximum size was 13 to 652,474 bp, with a median size of 8,909 bp (S1 Table). The incidence and scale of the LOH events vary according to the individual.

We validated the identification of L-SNV because there is a concern about false positive calls due to genotyping error, which can easily lead to violation of Mendelian inheritance at a random locus. To validate the identification of the L-SNV, we adopted a simulation-based approach to calculate a false positive rate. First, we arbitrarily choose a trio family that consists of a second-generation individual and one’s parents (first-generation). In this analysis, we used sample ID 4, which was identified with 64 L-SNVs that are comparable to the mean number of the identified L-SNV, and the individual’s parents (sample IDs 7 and 8). Then, we randomly marked 1% out of all variants of the three individuals as genotyping errors, respectively. Although it is known that the genotyping error rate of Genome Analysis Toolkit, which is a variant caller used in the CEPH/Utah cohort, is 0.1% ~ 1% [9], we adopted the 1% error rate to measure the false positive rate under stringent conditions. Finally, the false positive rate was calculated as the proportion of marked variants identified to be L-SNV. We carried out the above procedures 100 times repeatedly. The mean number and the standard deviation of the false positive calls of the sample ID 4’s L-SNV are 1.93 and 1.43, respectively. Hence, there can be about a 3.0% (1.93 / 64) false positive rate for the identification of the L-SNV in each individual. The false positive rate calculated here must be the maximum because the reported genotyping calling error rate of gatk is 0.1% ~ 1% [9].

### Relationship between LOH and parental age and sex

The emergence of postzygotic variants, which are genetic alterations that occur after fertilization, seems to be less affected by the condition of the parents compared with that of the variants that occur before fertilization [10–14]. For example, about 10% of postzygotic DNMs did not correlate with the parents’ sex and age, whereas about 90% of gonadal DNMs correlated with the parental parameters [7]. Therefore, we hypothesized that there is little correlation between the occurrence of LOH and parental sex and age, because the events occur between inter-homolog after fertilization.

To address this issue, first, we attempted to estimate the correlation between the incidence of LOH and parental age. Among the 132 first-generation individuals, the age range of the first-generation male at childbirth (second-generation individuals) was 18.4 (sample ID 147) to 47.2 (sample ID 389) years, whereas the age range of the first-generation female was 16.4 (sample ID 577) to 37.1 (sample ID 442) years. We found that there was no significant correlation between the incidence of LOH and paternal age (*r* = 0.061, *P* = 0.63, Fig 2A). A similar trend was observed for maternal age (*r* = −0.09, *P* = 0.47, S3 Fig). This result corroborates the findings of previous studies of postzygotic variation.

**Fig 2.**
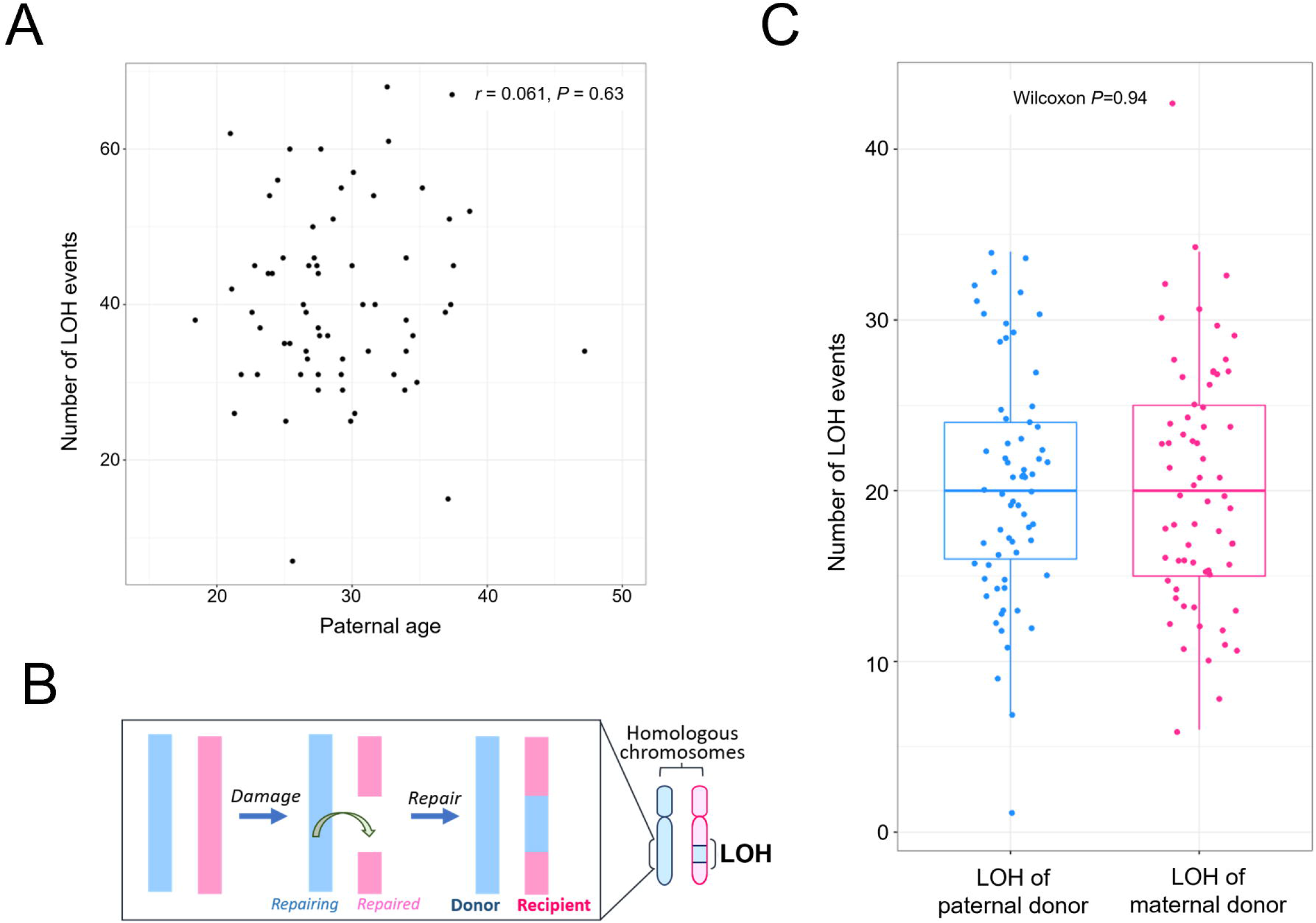
Relationships between LOH and parental age/sex. (A) Scatterplot between the number of LOH events and the paternal age (first-generation males). The relationship between the number of LOH events and the maternal age is presented in S3 Fig. (B) Conceptual diagram of the “donor” and “recipient” of homologous chromosomes. The segment in cyan indicates the part of the “donor” chromosome that repairs the damage of the “recipient.” The segment in magenta indicates the damaged part of the “recipient” chromosome that is repaired by the “donor.” (C) The boxplot indicates the number of LOH events that originated from paternal or maternal chromosomes.

To examine the effect of parental sex on LOH in terms of the mechanism underlying its occurrence, we classified the LOH region according to chromosome and discriminated their gamete of origin. In general, regarding LOH, the chromosome region at which a deletion occurred is termed “recipient”; concomitantly, the counterpart chromosome region that repairs the defect of the homolog is termed “donor” [15] (Fig 2B). We investigated the origin of the LOH region in the second-generation individuals. We observed that there was no significant difference between the number of sperm-originated LOH events and the number of egg-originated LOH events (Wilcoxon test, *P* = 0.94) (Fig 2C). This implies that the DNA lesion that gives rise to the LOH is not affected by a gamete bias. Taken together, these findings lead us to propose that the emergence of LOH is not affected by the parents’ age and sex, similar to that observed for postzygotic DNM.

### Distinction Between Gonosomal-mosaicism-associated LOH and Somatic-mosaicism-associated LOH

The postzygotic variants are distributed in various ways in the human body and sometimes affect not only the carrier individuals but also their progeny via transmission [10,12,13,16–18]. To examine the distribution of LOH events in germline and somatic cells, we investigated the transmission of the events to offspring (Fig 3A). Briefly, we set a 10-kbp window including heterozygote SNVs around the LOH events and confirmed the presence of the window in offspring. If the window including the recipient was observed in more than one offspring individual (third-generation individuals), it was considered a gonosomal-mosaicism-associated LOH, which is presented both in germline and somatic cells. In contrast, if the recipient before the occurrence of LOH, was similar to that of the progenitor (first-generation individual), and/or the donor window was observed, this was considered a somatic-mosaicism-associated LOH, which is present only in somatic cells [10,18].

**Fig 3.**
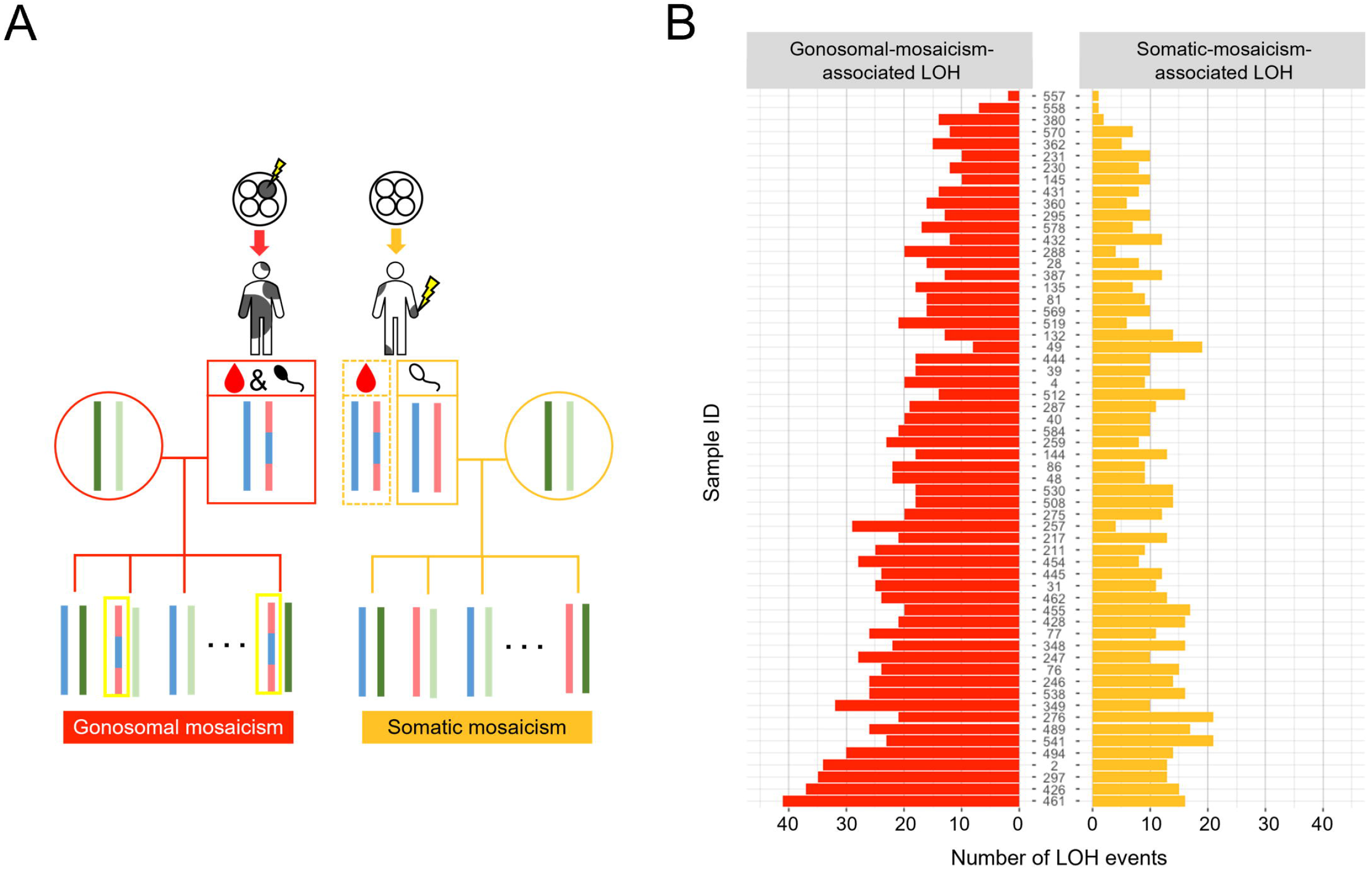
Distinction of gonosomal-mosaicism-associated LOH from somatic-mosaicism-associated LOH. (A) Schematic concept of gonosomal and somatic mosaicism. In the case of gonosomal mosaicism, the variant is present both in germline (black sperm icon) and somatic cells (blood icon) because the variant occurs at a very early embryonic stage. Therefore, LOH can be observed in one or more third-generation siblings (yellow rectangle). In the case of somatic mosaicism, the variant is not present in germline cells (white sperm icon). LOH was not observed in any of the third-generation siblings. (B) LOH was classified into gonosomal-mosaicism-associated LOH (red bar) and somatic-mosaicism-associated LOH (gold bar). The sample ID (at the center) is arranged in the order of increasing count, which is the sum of the two types of LOH.

We were able to track the transmission of 1,870 LOH events to offspring. There were 1,214 gonosomal-mosaicism-associated LOH events and 656 somatic-mosaicism-associated LOH events (Fig 3B). The average proportion of gonosomal vs. somatic mosaicism-associated LOH events was 0.65 vs. 0.35 among the individuals studied here. Notably, the incidence of gonosomal-mosaicism-associated LOH was approximately twice that of somatic-mosaicism-associated LOH in normal healthy individuals. It is probable that gonosomal-mosaicism-associated LOH, which can be observed in the blood cells of parents and offspring, occurs at a very early embryonic stage. Overall, given this variation in the inheritance mode of LOH among the individuals, we speculated that this yields different genome sequences among siblings and, hence, potentially has a deleterious effect on diseases such as tumorigenesis in individuals.

### Relationship between the occurrence of LOH and chromosomal features

The occurrence of HR repair during mitosis is typically associated with the degree of chromatin compactness and spatial distance of inter-homologs [19–22]. HR repair tends to be suppressed at heterochromatin more than it is at euchromatin, and the chance of repair increases at closer inter-homolog distances [20–22]. Similarly, regarding LOH events, we wondered whether they exhibit similar tendencies in light of the fact that these events are the result of HR repair after the defection of a single chromosome.

To address this question, we measured the GC content of the genomic region including the LOH. The GC content of this region has been shown to strongly correlate with chromatin compactness [23]. We first investigated the enrichment of all 2,684 LOH events by comparing the frequency of LOH and GC content on the reference genome fraction, which was cleaved at 1 kbp as a window. At all LOH events, the enrichment of the events was likely to increase as the GC content increased, from about 45% (Fig 4A). In particular, we observed that LOH tended to be enriched in GC-rich windows, from around >60% to ≤65%. This result corroborates the assumption that HR repair seems to be suppressed at heterochromatin in the genome.

**Fig 4.**
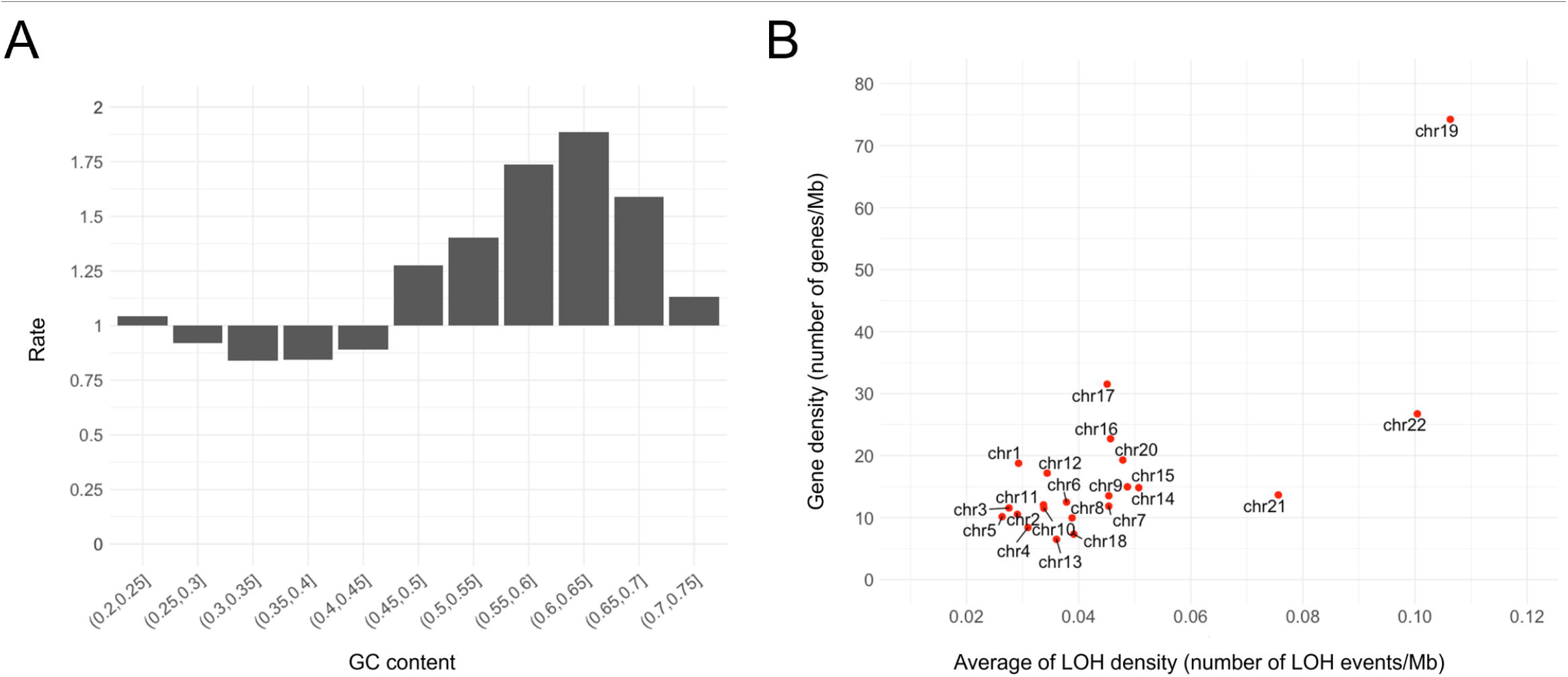
Relationship between the occurrence of LOH and chromosomal features. (A) Histogram of LOH enrichment according to GC content. The bar indicates the ratio of the observed LOH to the expected LOH according to the GC content of the genome regions. A ratio <1 means that LOH events are less enriched at that GC content, whereas a ratio >1 means that LOH events are enriched at that GC content. (B) Scatterplot of the gene density per chromosome vs. the average number of LOH events per chromosome.

In the human nucleus, gene-rich chromosomes, such as chromosome 19, are localized in the internal part of the nucleus, whereas gene-poor chromosomes, such as chromosome 18, are positioned at the periphery of the nucleus, near the nuclear lamina [22,24–28]. If chromosomes are located in the internal part of the nucleus, they become spatially closer than if they are located at the periphery of the nucleus; accordingly, the inter-homolog distances are reduced and the chance of HR repair increases [22]. Therefore, we hypothesized that LOH occurs at gene-rich chromosomes more frequently than it does at gene-poor chromosomes because of the shorter inter-homolog distances.

Here, we observed that chromosome 19 was strikingly high in both gene density and mean number of LOH events, whereas chromosome 18 was low in both features (Fig 4B). It is well known that although inter-homolog distances are usually larger than inter-heterolog distances, the inter-homolog distance of chromosome 19 is small [24–26,28]. Chromosomes 1, 4, 10, 14, 16, and 18 seem to reflect the fact that the inter-homolog distances are larger than the inter-heterolog distances [22]. Chromosomes 21 and 22 exhibited a relatively high mean number of LOH events, even though their gene density is lower than 30 genes/Mb. This result seems to confirm the results of a previous study that showed that shorter chromosomes tend to exist in the interior of the nucleus [25,27,29–31]. In particular, chromosome 21 is preferentially situated in the internal region of the nucleus, because this chromosome is acrocentric and includes nucleolar organizer regions [22,30,32,33]. Taken together, these results suggest that the chance of the emergence of LOH depends on the gene density associated with chromatin compactness and inter-homolog distances, as illustrated by the radial position of a chromosome associated with CT.

## Discussion

We quantitatively identified LOH events in germline as well as somatic cells by taking advantage of the SNV data of three-generation families. We identified approximately 40.7 LOH events per individual. This number is comparable to the number of DNMs per individual, i.e., 70, reported previously [7]. Intriguingly, more than half of the LOH events exhibited gonosomal mosaicism. Genome variants tend to be patrolled and/or repaired more strictly in germline cells, as they are transferable to progeny, compared with in disposable somatic cells [34]. The fact that the DNM rate in germline cells is lower than that detected in disposable somatic cells is a representative example of such a phenomenon [35,36]. In contrast, considering that LOH results from DNA repair after genome defection, it is possible to infer that the incidence of LOH exhibits an opposite trend to that observed for DNM. Furthermore, we revealed the possible association between CT and the occurrence of LOH by investigating the genomic features of the LOH regions. Specifically, the incidence of LOH is inversely proportional to the inter-homolog distances, which depend on several parameters, such as gene density and chromosome length.

Two issues should be addressed in the interpretation of our findings. The first point is that the number of LOH events identified in this study may have been underestimated for the following two reasons. (1) We applied stringent filtering to remove repeat regions, unassembled regions, and LCR. In fact, studies of genetic alterations have mentioned that HR spontaneously occurs in repeat regions [37]. Nevertheless, to ensure the accuracy of our results, we excluded such regions, because they may have a low sequencing quality. (2) The raw data used here were sequenced from peripheral blood cells exclusively; i.e., tissue-specific LOH events occurring in tissues other than blood were not included in our results. For example, for the onset of myeloproliferative disorders, the LOH has to occur in T cells [2]. Therefore, we suggest that the LOH identified in this study may have been sufficiently frequent for detection by occurring in hematopoietic stem cells or in very early embryonic stem cells.

The second point is that the proportion of LOH associated with gonosomal mosaicism observed in this study may actually be higher in reality. In this study, LOH was classified into two groups, i.e., gonosomal-mosaicism-associated LOH and somatic-mosaicism-associated LOH, by investigating whether the LOH was transmitted to offspring. The number of progenies is particularly important in this regard because it is directly related to the accuracy of the results [12,38,39]. If, for instance, the third-generation includes 4 individuals, the probability that the LOH that occurred in the parent is not transmitted to any offspring is (1 / 2) ^ 4, which is not negligible. Although human beings inherently do not have many offspring compared with other organisms, the Utah population used here, fortunately, has a relatively high birth rate because of their religious beliefs. Therefore, we included only families that had 7 or more third-generation individuals in our study, to take full advantage of the characteristics of the cohort [average of about 8.97 children (third-generation) per family].

Inherited variants in tumor suppressor genes are thought to be the main cause of hereditary cancer [40,41]. In some tumors, the inherited pattern causes tumors more frequently than does the sporadic pattern. For example, in some cases, hematologic malignancies are incurred from the duplication of the mutated allele in specific gene such as *JAK2* gene via LOH [5]. Although cancers with a hereditary susceptibility represent 5%–10% of all types of cancer [41], the nature of LOH that is the trigger of the disease as the second hit has received little attention. We expect that the findings pertaining to the incidence of LOH obtained in this study will provide clues to infer the onset rate of cancer among families with a hereditary cancer susceptibility.

## Materials and Methods

### Dataset

VCF files were downloaded from the National Center for Biotechnology Information (NCBI)’s dbGaP, and the dataset title was “Genome sequencing of large, multigenerational CEPH/Utah families” (https://www.ncbi.nlm.nih.gov/projects/gap/cgi-bin/study.cgi?study_id=phs001872.v1.p1). The dataset consisted of 603 individuals from 33 large families in Utah in the United States. The families comprise three biological generations, including offspring, parents, and grandparents. Pedigree information of these families was also obtained from the same study. Here, we used 285 individuals from 33 three-generation immediate families, which are a subset of the 33 large families and met the following conditions: (1) an intact family with usable SNV data for all family members (S1 Fig), and (2) presence of 7 or more siblings in the third-generation of a family.

### Identification of L-SNVs and LOH

First, we performed the following quality control of the VCF files to identify L-SNVs that were considered to be included in the LOH. The variant had to satisfy the condition of GATK HaplotypeCaller [42] as “PASS”, a read depth ≥12, and a Phred-scaled genotype quality ≥20[39]. The DNA sequences that corresponded to repeat regions and LCR were excluded [the data were downloaded from RepeatMasker [43] (Genome Reference Consortium Human Build 37; GRCh37) (http://www.repeatmasker.org) of the UCSC genome browser [44] and https://raw.githubusercontent.com/lh3/varcmp/master/scripts/LCR-hs37d5.bed.gz [45,46], respectively]. We only used variants located in autosomes to avoid a bias originating from sex.

An L-SNV was defined as a homozygous locus with a genotype originating from a single parent that did not exist in combinations of parents, as assessed by referring to the genotype fields of the VCF format (S4 Fig). For example, if parents’ genotypes are 1/2 and 2/2 and the progeny’s genotype is 1/1, the progeny’s SNV is regarded as an L-SNV. If, however, the progeny’s genotype is 2/2, the SNV is excluded as a normal homozygous variant because the SNV is included in the possible combinations from the parent’s genotype. If the progeny’s genotype is 3/3, this SNV is excluded as a de novo variant or a variant call error. This is because any parents do not have allele 3, although the genotype is not included in the possible combinations from the parents’ genotype. To discriminate between the homologous “recipient” region, in which the LOH literally occurred, and the “donor” region, which acts as a template for DNA repair from L-SNV, we phased the filtered SNVs to comply with Mendelian inheritance using Beagle version 4.0 [47]. We then phased the L-SNVs manually because of their relatively low phasing accuracy, which is attributable to the characteristics of Mendelian inheritance errors. Finally, the LOH was inferred from the phased L-SNVs and SNVs. We defined the LOH regions as consecutive homologous regions including L-SNVs and optionally homozygous SNVs that were restricted by the nearest heterozygous SNVs on both sides (S2 Fig).

### Assessment of the effect of parental age and sex on the occurrence of LOH

To assess the effect of parental age on the occurrence of LOH in offspring, we investigated the correlation between the incidence of LOH in second-generation individuals and the age of the first-generation individuals. We obtained the information of the age and sex of first-generation individuals from https://github.com/quinlan-lab/ceph-dnm-manuscript. The parental age was rounded off to one decimal place in this study. Correlations with a *P*-value < 0.5 (Pearson’s coefficient) between the incidence of LOH and parental age were estimated using the default option of the “ggpubf’ package (v. 0.4.0) (https://rpkgs.datanovia.com/ggpubr/index.html) of the R (v.4.0.2) (The R Project for Statistical Computing, Vienna, Austria) software and visualized using the same package. The same approach was employed to assess and visualize the effect of parental sex on LOH. In this case, the Wilcoxon test was used to compare the mean number of LOH events.

### Discrimination between gonosomal-mosaicism-associated LOH and somatic-mosaicism-associated LOH

To discriminate between LOH associated with gonosomal mosaicism and that associated with somatic mosaicism, we tracked the mode of transmission of the LOH to the offspring. For this analysis, we used the 10-kbp window surrounding the LOH that contained the heterozygous as well as the homozygous variants, such as the L-SNV. If the “recipient” window was not observed in any of the offspring, the LOH in the window was considered to be associated with somatic mosaicism, which is only present in somatic cells. In contrast, if the “recipient” window was observed in one or more siblings, the LOH was considered to be associated with gonosomal mosaicism, which is present in both germline and somatic cells. During the comparison of the window of second- and third-generation individuals, the variants existing in the window had to completely match each other.

### Estimation of GC content and gene density

We downloaded the information pertaining to chromosome length from NCBI’s (https://www.ncbi.nlm.nih.gov/assembly/GCF_000001405.13/) GRCh37. The GC content was calculated using the following formula.

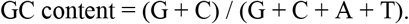

Where A, C, G, and T indicate the number of each nucleotide. During calculation of GC content, we only used a 1-kbp window that contains more than 100 nt after removing the LCR, repeat region, and unassembled region. To measure gene density, we counted the number of “protein_coding” genes at each chromosome, followed by calculations using the filtered chromosomal length. We used the comprehensive gene annotation of the GENCODE project (https://www.gencodegenes.org/human/release_19.html, GRCh37.p13).

## Acknowledgments

This work was supported by JST SPRING, Grant Number JPMJSP2136.

## Supporting information

S1 Fig. Example of division of a large family.

S2 Fig. Schematic concept of minimum and maximum size of LOH.

S3 Fig. Relationship between LOH and maternal age.

S4 Fig. Flow chart of L-SNV identification.

S1 Table. List of loss of heterozygosity identified in this study.

## Data reporting

### Data availability

The whole-genome sequencing datasets utilized in this study are available in NCBI Sequence Read Archive (SRA) and dbGaP with the accession number phs001872.v1.p1. All source data generated in this study are available at https://github.com/rgwluj123/LOH_3generation_Utah.

### Code availability

The code developed for this study are freely available at https://github.com/rgwluj123/LOH_3generation_Utah.

