## Supplemental Figures for "Genome-wide identification of loss of heterozygosity generated by mitotic homologous recombination reveals its possible association with spatial positioning of chromosomes"

**For**

A

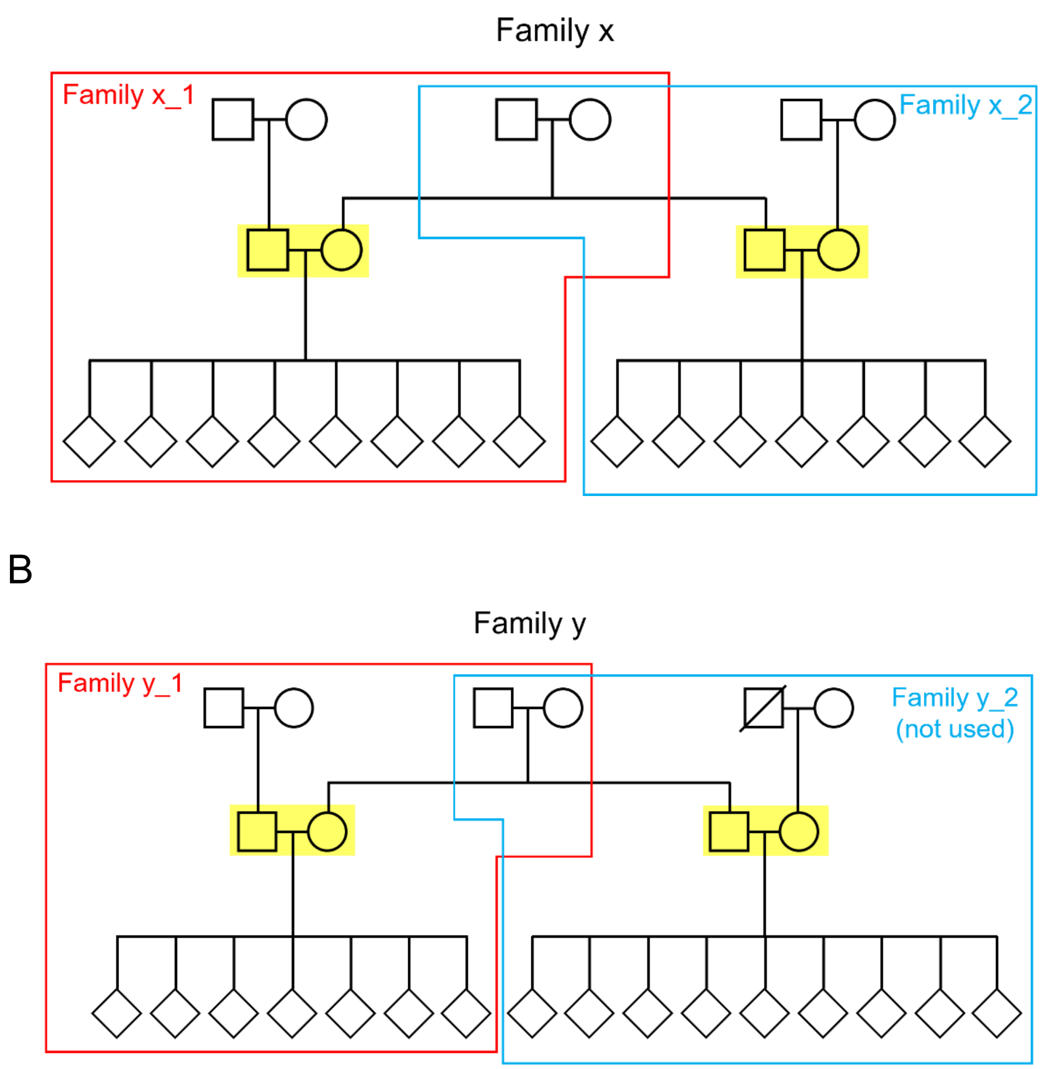


S1 Fig. Example of division of a large family. Immediate families used in this study were selected based on the married couples in the second-generation in a large family. (A) Family x separates into family x_1 (box in red) and family x_2 (box in cyan) by the married couples in the second-generation (box in yellow). (B) Family y separates into family y_1 (box in red) and family y_2 (box in cyan) by the married couples in the second-generation (box in cyan). However, the family y_2 would be excluded in this study because of the lack of data on member (slashed individual).

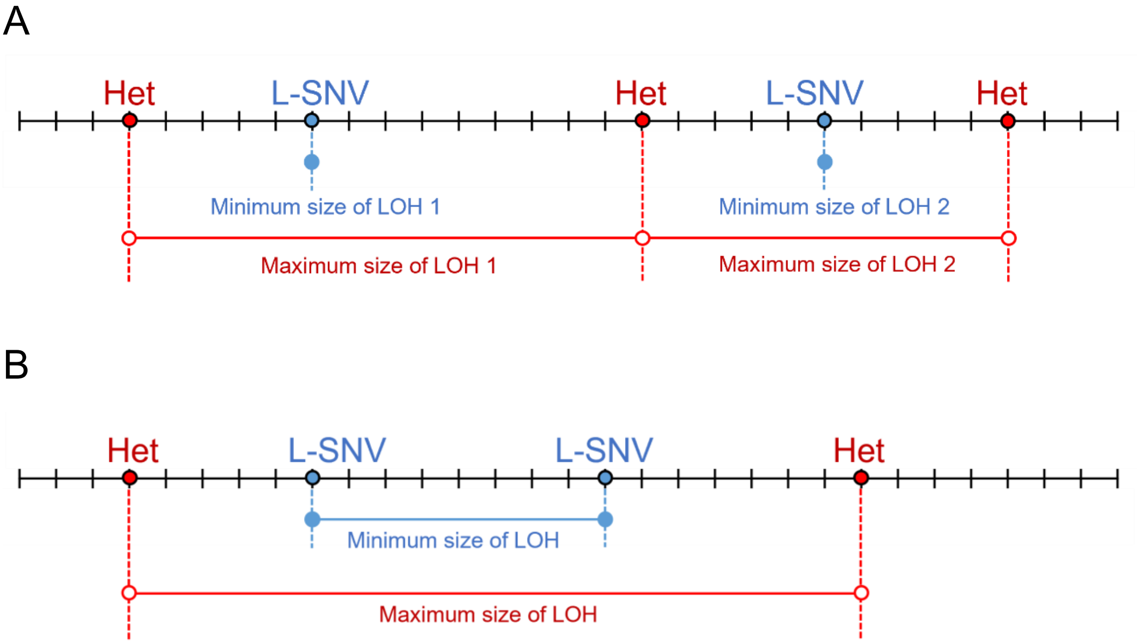


S2 Fig. Schematic concept of minimum and maximum size of LOH. The minimum size represents the distance from the first L-SNV to the last L-SNV in an LOH event. The maximum size represents the distance between two nucleotides just before the first heterozygous variant upstream and downstream from an LOH event. (A) This figure shows the case of one L-SNV between heterozygous variants. (B) This figure shows the case of two or more L-SNVs between heterozygous variants. Het, heterozygous variant

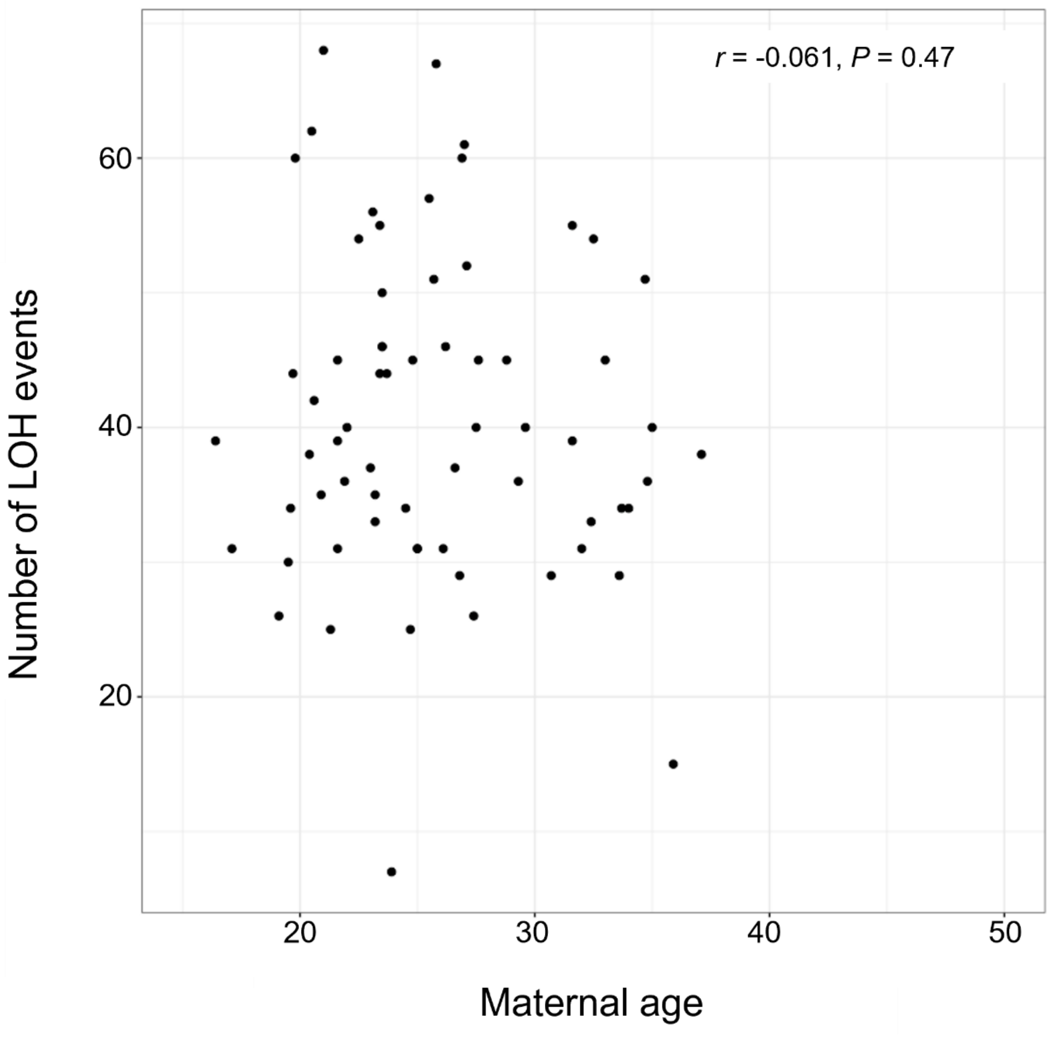


S3 Fig. Relationship between LOH and maternal age. Scatterplot between the number of LOH events and the maternal age (first-generation females).

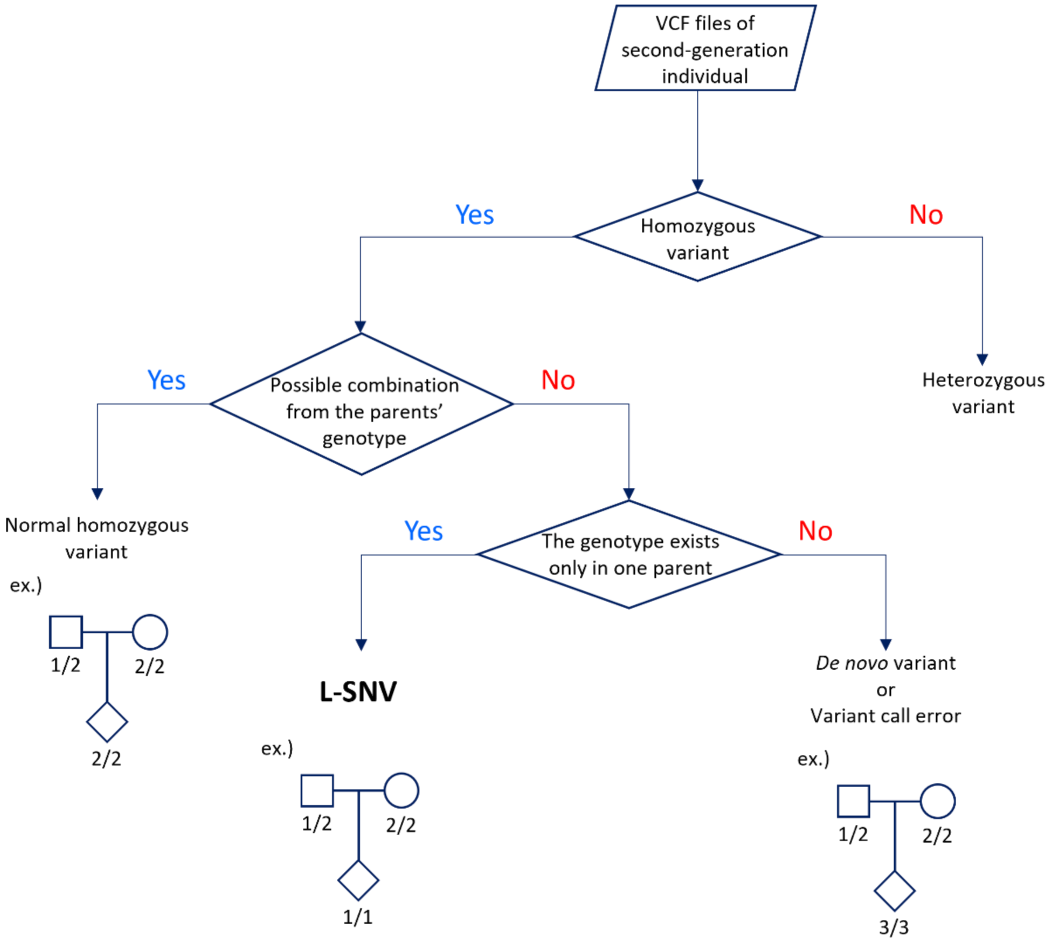


S4 Fig. Flow chart of L-SNV identification. The flow chart illustrates the pipeline of the L-SNV identification constructed in this study. Pedigrees below the flow chart present examples of the variants.
